# Sources of predictive information in dynamical neural networks

**DOI:** 10.1101/2019.12.23.887554

**Authors:** Madhavun Candadai, Eduardo J. Izquierdo

## Abstract

Behavior involves the ongoing interaction between an organism and its environment. One of the prevailing theories of adaptive behavior is that organisms are constantly making predictions about their future environmental stimuli. However, how they acquire that predictive information is still poorly understood. Two complementary mechanisms have been proposed: predictions are generated from an agent’s internal model of the world or predictions are extracted directly from the environmental stimulus. In this work, we demonstrate that predictive information, measured using mutual information, cannot distinguish between these two kinds of systems. Furthermore, we show that predictive information cannot distinguish between organisms that are adapted to their environments and random dynamical systems exposed to the same environment. To understand the role of predictive information in adaptive behavior, we need to be able to identify where it is generated. To do this, we decompose information transfer across the different components of the organism-environment system and track the flow of information in the system over time. To validate the proposed framework, we examined it on a set of computational models of idealized agent-environment systems. Analysis of the systems revealed three key insights. First, predictive information, when sourced from the environment, can be reflected in any agent irrespective of its ability to perform a task. Second, predictive information, when sourced from the nervous system, requires special dynamics acquired during the process of adapting to the environment. Third, the magnitude of predictive information in a system can be different for the same task if the environmental structure changes.

**Significance Statement:** An organism’s ability to predict the consequences of its actions on future stimuli is considered a strong indicator of its environmental adaptation. However, in highly structured natural environments, to what extent does an agent have to develop specialized mechanisms to generate predictions? To study this, we present an information theoretic framework to infer the source of predictive information in an organism: extrinsically from the environment or intrinsically from the agent. We find that predictive information extracted from the environment can be reflected in any agent and is therefore not a good indicator of behavioral performance. Studying the flow of predictive information over time across the organism-environment system enables us to better understand its role in behavior.

---

**P**redictive coding is emerging as a strong candidate for its ability to provide a general framework for understanding the neural basis of behavior (1–4). The idea is that organisms encode information about future environmental stimuli in their neural activity based on their knowledge of the environment. Intuitively, an organism that is able to predict the consequences of its action on its future sensory experiences is more likely to be adapted to its environment. There are two prominent research fronts that study the role of predictive coding in behavior: the hierarchical predictive processing framework (5, 6) and the efficient coding principle (7, 8). These two fronts are complementary because they address different aspects of how a nervous system acquires predictive information. The hierarchical predictive processing framework focuses on how predictions are generated in the organism’s brain. The efficient coding principle focuses on how the nervous system extracts predictive information from environmental stimuli. Both theories have been supported by experimental evidence, primarily in the visual and auditory systems (9–12).

In living organisms, predictive information is likely acquired from a dynamically changing contribution of the environment and the agent’s own internal dynamics (2). Consequently, although different systems may be equally predictive about their future stimuli, the operation of their nervous systems may be entirely different. Therefore, understanding the role of predictive information in behavior requires that the source of information is identified. In this paper, we address the following questions. How do we identify the source of predictive information and study its dynamics during a behavior? Does tracking the source of predictive information better explain an agent’s ability to perform a task? What are the factors that influence the source and magnitude of predictive information encoded in a neural network?

In the first part of this paper, we demonstrate that predictive information will generate indistinguishable results for systems that are at the two extremes of potential agent-environment interaction: a system whose only source of predictive information is the nervous system and a system whose only source of predictive information is the environmental stimuli. In order to better understand how the nervous system generates predictive information, we propose that it is essential to decompose information transfer across the different components of the system and to track the flow of information in the agent-environment system over time. The principal contribution of this paper is an information-theoretic framework to quantify the contributions from the nervous system and the contributions from the environmental stimuli to the total predictive information in an agent. First, we decompose the total predictive information in the neural system into information that was uniquely transferred from each source. In order to do this, we employ multivariate extensions to information theory (13). Second, we unroll information over time to backtrack the origin of the source of predictive information and how they change over time. To validate the proposed theoretical framework, we examine it on a set of computational models of agent-environment systems, where the agent is driven by a dynamical recurrent neural network (14, 15). The systems have been deliberately designed so that the source of predictive information is tractable and manipulable. We demonstrate how the proposed framework correctly reveals different sources of predictive information in systems with otherwise similar amounts of predictive information. Ultimately, we demonstrate how revealing the flow of information across the agent-environment system can help us to better understand the mechanisms underlying predictive coding.

Predictive information is studied in living organisms because it is considered a signature of their adaptive capacities (5, 8, 9). In the second part of this paper we study the relationship between a system’s ability to perform a task and its predictive information. In order to do this, we turn to a computational model of an agent that is required to process the received stimulus from the environment and make a decision based on it. Specifically, we study predictive information in the context of a relational categorization task (16, 17). We generate model systems that are adapted to their environment and yet remain tractable to analysis by optimizing dynamical recurrent neural networks using an evolutionary algorithm to perform the task (18, 19). We then proceed to analyze the resulting systems using predictive information and we compare the results against that of random systems that cannot solve the task. Counterintuitively, we observe that predictive information in trained neural networks is similar to predictive information in random neural networks. This suggests that predictive information alone is not sufficient to distinguish between living organisms that are adapted to their environments and non-adaptive systems. The rest of the paper focuses on an analysis of optimized and random systems using the framework proposed. Altogether, we demonstrate that decomposing predictive information across the components of an agent-environment system, and unrolling it over time reveals its true nature.

## Identifying the source of predictive information

Predictive information is the information encoded in neural activity about its future stimulus. Formally, it is defined as mutual information between current neural activity (*N*_*t*_) and the stimulus at a future time (*S*_*t*′_) (9, 20–23), according to:

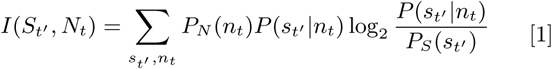

where *t*′ = *t* + *δt* with *δt* > 0, *P*_*S*_ is the distribution of environmental stimuli, *P*_*N*_ is the distribution of neural activity across the entire experiment, *P* (*s*_*t′*_ |*n*_*t*_) is the conditional probability that the stimulus is *s* at a future time *t*′ given that we have observed a neural activity of *n* at time *t*. When this measure is estimated using the stimulus and neural activity across all data points separated in time by some *δt*, it is a measure of reduction in uncertainty in future stimulus given the current neural activity.

The presence of predictive information in a neural network suggests there is a source where this information was generated. In an idealized agent-environment system (Fig. 1A), the source of information can be either the neural activity in the previous time step, the environmental stimulus in the previous time step, or both (Fig. 1B). Measuring predictive information as defined in equation 1 requires that we examine two variables: current neural activity (*N*_*t*_, henceforth *X*) and future stimulus (*S*_*t*+*δt*_, henceforth *Y*). Identifying the source of this predictive information requires that we examine two additional variables: past neural network activity (*N*_*t*−*δt*_, henceforth *A*) and past stimulus (*S*_*t*−*δt*_, henceforth *B*). Such an analysis requires that we adopt multivariate extensions to information theory. We focus specifically on Partial Information Decomposition (PID) (13), a method for decomposing multivariate mutual information into combinations of unique, redundant and synergistic contributions, as well as measures of information gain, loss and transfer (13, 24–32). In order to identify the source of predictive information, we can decompose the total information that the current neural activity has about the future stimulus into three components: (a) information uniquely transferred from past environmental stimulus, *T*_*Y* ;*A*→*X*_ ; (b) information uniquely transferred from past neural network activity, *T*_*Y* ;*B*→*X*_ ; and (c) information redundantly transferred from past environment stimulus and past neural network activity, *T*_*Y* ;*{A,B}*→*X*_, according to:

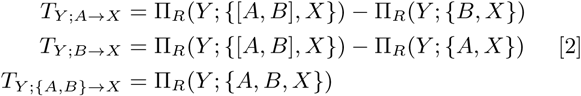

where Π_*R*_(*Y* ; {*A*_*1*_, *A*_*2*_, ..*A*_*3*_}) is the redundant information that random variables *A*_1_ through *A*_*k*_ have about the random variable *Y* and [*A, B*] refers to a random variable that is a concatenation of *A* and *B*. In words, information about *Y* transferred uniquely from source *A* to *X* is estimated as the total redundant information from the combined sources [*A, B*] *minus* the information that is redundant with the other source *B*. This decomposition of the total information into different contributions is typically represented using a PI-decomposition diagram (Fig. 1C). Several approaches have been proposed to measure redundant information, Π_*R*_ (24, 33, 34). Here, we use *I*_*min*_ because this is the only approach that can guarantee non-negative information decomposition in a system with four random variables, as is the case here.

**Fig. 1.**
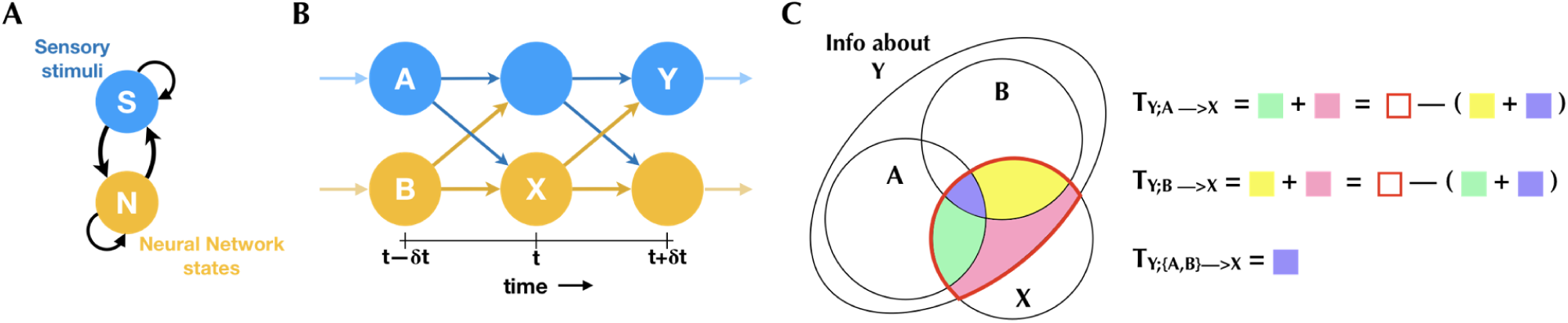
Predictive information source estimation based on idealized agent-environment interaction. [A] Sensory stimuli (S) and neural activity (N) are two coupled dynamical systems. [B] Agent-environment interaction unrolled over time. *X* represents current neural activity, *N* (*t*), *Y*, future environmental state, *S*(*t* + *δt*), and *A* and *B* represent the sources, namely past neural activity *N* (*t* − *δt*) and past environmental state, *s*(*t* − *δt*) respectively. [C] Partial information diagram for calculating the sources of predictive information in an agent-environment system. The total information that *X* has about *Y* is a combination of information that is available uniquely from *A* alone (green), uniquely from *B* alone (yellow), synergistically from their combination [*A, B*] (pink), and redundantly from both of them (purple). PID allows us to measure information transfer using these components. Alternatively, they can also be measured by estimating the total redundant information from both sources combined (red) and removing the information from the other source.

During the course of behavior, the flow of information in a system changes over time (35, 36). In order to understand the source of predictive information for any agent-environment system, it is not enough to decompose information from multiple sources; we must also track its flow of information over time. Although information theoretic measures are typically averaged over time, the measures described above can be unrolled over time (36, 37). This is done by measuring information transfer at each time-point using data collected across several trials thereby allowing us to study the dynamics of predictive information sources.

## Disparate systems with similar predictive information

Neural systems can be predictive in fundamentally different ways: they can generate predictive information internally or they can extract it from environmental stimulus. We use computational models of two extreme conditions where the ground-truth predictive information source is known to be the environment in one condition and the neural network in the other, to demonstrate that (a) predictive information cannot distinguish between these different kinds of systems and (b) it is only through decomposing the information across sources and unrolling over time that we can distinguish the two systems based on their operation. The two conditions we consider are agent-environment interactions at two extremes of the range of possible interactions: a central pattern generator (CPG) and a passive perceiver (PP). In the CPG condition, the neural network influences the environment by producing spontaneous oscillatory activity but receives no input from the environment (Fig. 2A). In the PP condition, the neural network is influenced by input from the environment, but it does not affect the environment (Fig. 2B). We evolved 100 different dynamical recurrent neural network CPGs, and in each case, we fed the sum of the neurons’ outputs to the environment (Fig. S1A,B). For the PPs, we generated 100 random neural networks and fed them an oscillatory input. In order to provide the same distribution of activity as the CPG condition, we provided the random neural networks with the same oscillatory environmental signal that CPGs generated (Fig. S1C). The environmental signal and neural data were recorded from each instance for 500 trials where, in each trial the environment started with a different initial condition. Although, the environmental signal and the neural activity exhibit oscillatory activity in both conditions, the key difference in the operation of these systems is that in the CPGs the neural network drives its own activity and in the PP, the environment drives the neural network. Therefore, the neural network is the source of predictive information in the CPGs and the environment is the source of predictive information in the PPs.

**Fig. 2.**
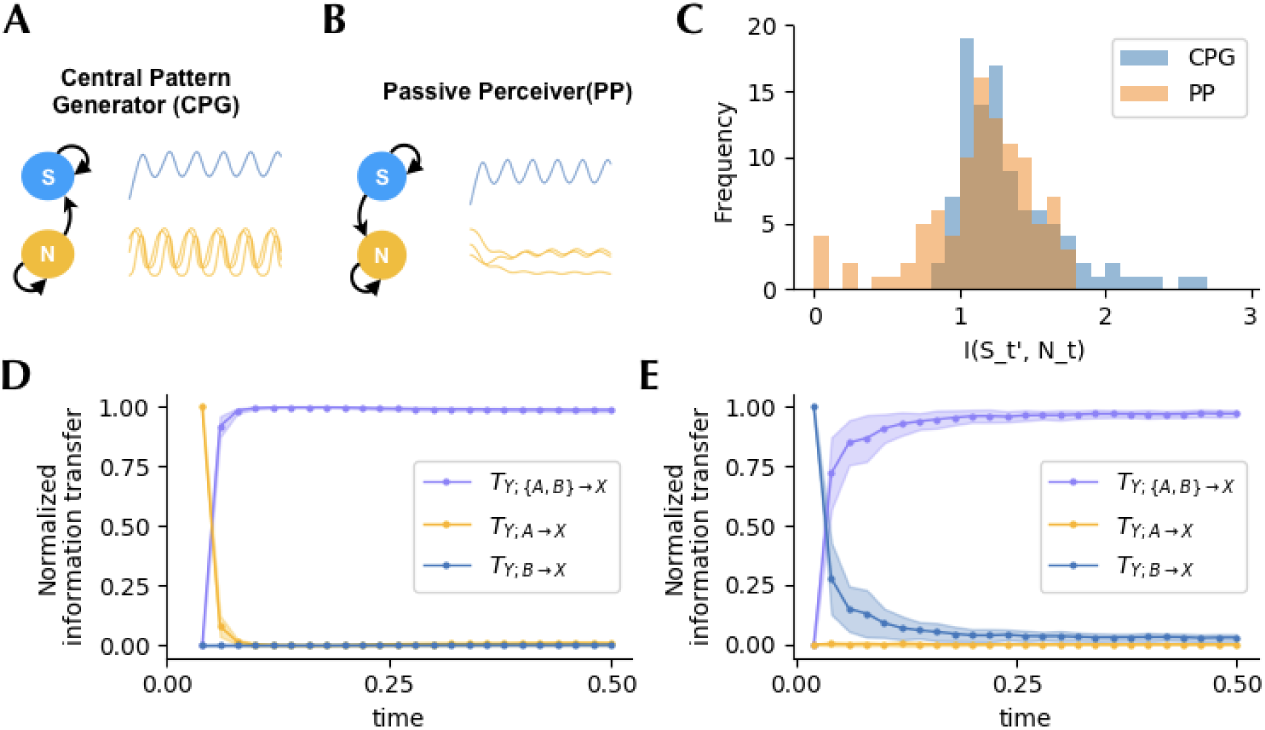
Predictive information in systems on the extremes of the range of possible agent-environment interactions [A] Schematic and traces of a Central Pattern Generator (CPG) that influences the environment through intrinsically generated oscillations. [B] Schematic and traces of a Passive Perceiver (PP) that is driven by oscillatory inputs from the environment (in this case, by the environmental signals recorded from the CPGs) [C] Estimating total predictive information as shown in equation 1 shows that CPG and PP models encode similar amounts of predictive information about environmental state in the next time-step. [D] Decomposing that total information into information that came from the environment and the neural network consistently showed that information about the next time-step in the CPG originated in the neural network (yellow) before becoming redundant (purple) as the environment and the neural network synchronized. [E] Conversely, with PPs, the environment was consistently shown to be the source of information (blue) before they environment and neural network synchronize and become redundant (purple).

As a first step in the analysis of these two systems, we used the recorded data to measure predictive information in the neural network about the environmental signal in the next time-step (*δt* = 0.02*s*). To calculate predictive information, data distributions were constructed using all tuples of neural activity at time *t* and environmental signal at time *t* + *δt*, averaged across time and trials. The analysis revealed that the neural networks, in these two otherwise diametrically opposed systems, encoded similar levels of information about stimulus in the next time step (Fig. 2C). From this first experiment, we conclude that predictive information is not sufficient to distinguish systems that generate their own predictive information from systems that encode the information available from the environmental stimuli.

To understand what makes these two neural systems different, it is necessary to identify the source of their predictive information. As a next step in our analysis, we decomposed the information in the neural system about the future stimuli across the different possible sources and we unrolled the analysis over time. At each time-point, we measured information in the neural network about the environmental signal in the next time-step that was uniquely transferred from the environment, uniquely transferred from the neural network and redundantly from both.

In the CPG condition, since the neural networks are not influenced by the environment (Fig. 2A), the only source of information about the future environmental signal is from the neural network itself. Accordingly, the dynamics of information transfer for CPG systems reveals correctly that the neural network is the source of predictive information (Fig. 2D). At the start of the interaction between agent and environment, the neural network uniquely transfers information about the future environmental state to the environment. Following that, the environment quickly becomes synchronized with the neural activity. This means that the state of the environment becomes informative of its own future state. This results in the environment and the neural network becoming redundant sources of predictive information. Crucially, however, the environment never provides any unique information to the neural network about its future stimulus.

In the PP condition, since the neural networks are driven by the environment (Fig. 2B), the only source of information about the future environmental signal is the stimulus from the environment itself. Accordingly, the dynamics of information transfer for PP systems reveals correctly that the environment is the source of predictive information (Fig. 2E). As opposed to the CPG systems, at the start of the interaction between the neural network and the environment, it is the environment that transfers unique information to the neural network. Subsequently, and similarly to the CPG condition, as the state of the neural network begins to encode the information from the environmental stimulus, the predictive information is redundantly transferred by both the neural network and the environmental stimulus. Consistent with our expectation, the neural network never provides any unique information to itself about the future of the stimulus.

In summary, in this section we show that predictive information alone cannot distinguish between two extremely different kinds of neural systems, both of which encode predictive information about the future of the environment. This is because when the entire time course of the data is considered, the environment and neural network are synchronized for a majority of the time. Information uniquely transferred from any source is only detectable within a short time window before they synchronize. In this section, we have shown that decomposing information across sources and unrolling over time allows us to study information source dynamics at every perturbation to the agent-environment interaction and hence reveals the source of predictive information.

## Predictive information with structured stimuli

The natural environment is not uniformly random but is in fact highly structured with spatial and temporal regularities (2, 38, 39). This structure is reflected in the stimulus that agents receive from the environment. Accordingly, this is emulated in most preparations in neuroscience, where a neural system is presented with artificial stimuli with some underlying structure designed by the experimenter. We posit that the structure in the environment will strongly influence the amount of predictive information encoded by the neural network and its sources. In order to study this, we examined the flow of information in a neural network model trained to solve a relational categorization task.

Relational categorization is the ability to discriminate objects based on the relative value of their attributes (16, 17). This task allows us to specify the inherent structure in the environment by changing the distribution of objects whose attributes are compared thus making it especially suited for studying the influence of environmental structure on predictive information. It involves providing the neural network with stimuli across three stages: cue, delay, and probe. In the cue stage, the neural network is provided with a stimulus of specific magnitude for a duration of time. This is followed by a delay stage, where no stimulus is provided. Finally, in the probe stage, the neural network is provided with a second stimulus. The magnitudes for the cue and probe stage stimuli are picked from a predesignated distribution (Fig. 3A). It is this distribution that defines the structure in the environment. For this study, we design it such that the stimulus in the probe stage can have a magnitude that is one of two values: smaller (*cue* − 1) or larger (*cue* + 1) than the stimulus provided during the cue stage (Fig. 3B). The goal of the neural network in this task to perform a relational categorization of “greater than” or “lesser than” by producing an output of +1 or − 1 respectively, during the probe phase. This task has been widely studied in a variety of contexts including in humans (40), pigeons (41), rats (42), insects (43), as well as using computational models (44, 45).

**Fig. 3.**
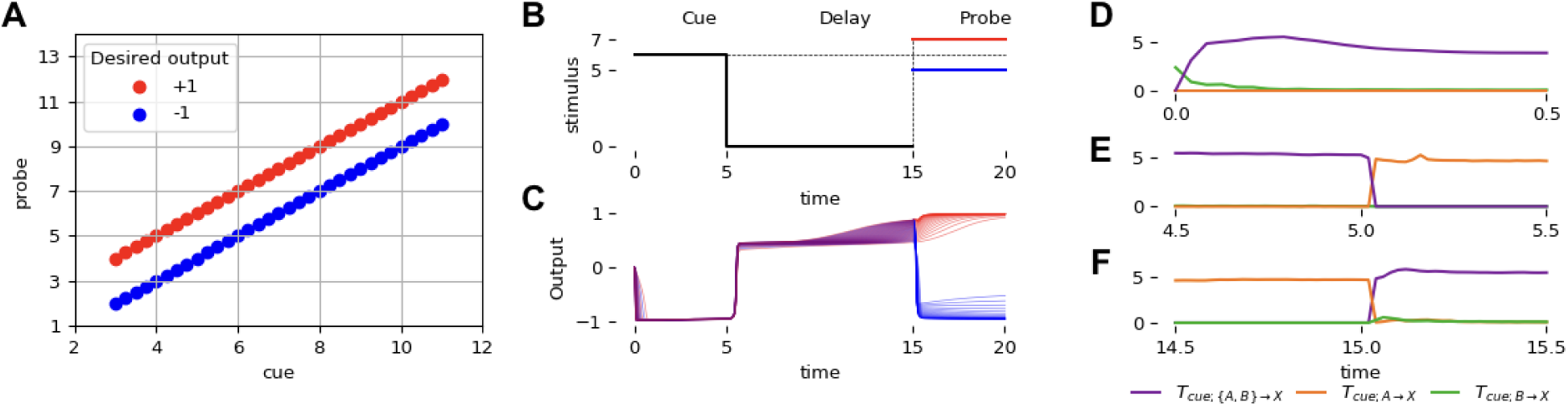
Predictive information source dynamics with structured stimuli. [A] Distribution of cue and corresponding probes in the relational categorization task. For each cue, the probe can be one of two values: greater, *cue* + 1, or lesser, *cue* − 1, with the expected outputs of +1 (red) and −1 (blue) respectively. [B] One trial of the relational categorization task. The cue stimulus is presented till t=5, followed by a delay period with no stimulus (t=5 to t=15) and then a probe that is greater (red) or lesser (blue) than the cue is provided. [C] Behavior of the best out of 100 dynamical neural networks optimized to perform this task showing perfect categorization of the relational value from 35 trials where the probe was greater (red) and 35 where the probe was lesser (blue). [D] Dynamics of information about the cue during the cue stage show information uniquely provided by the environment (green) initially, but becoming redundantly available in the neural network and environment (purple) as it encoded the cue. [E] Towards the end of the cue stage, information is entirely redundant (purple). When the stimulus stops being provided at t=5, the neural network is the unique source of information about the cue (orange). [F] Dynamics of information about the cue just before the probe arrives showing that the neural network continues to retain information about the cue (orange). At t=15, when the probe is provided, information quickly becomes redundant (purple) denoting that the probe has information about the cue.

In this section, we show results from analysis of neural networks performing the relational categorization task. We demonstrate that decomposing information across the sources and unrolling over time reveals that the environment is structured by appropriately attributing the observed predictive information to either the environment or the dynamics of the neural network. Furthermore, we demonstrate that encoding predictive information alone is not indicative of task performance and that the magnitude and source of predictive information can change during the course of a behavior depending on environmental structure and neural network dynamics.

### Characterizing information source dynamics in the best optimized neural network

Dynamical recurrent neural networks were optimized using an evolutionary algorithm to perform the relational categorization task. A total of 100 independent evolutionary runs yielded an ensemble of 100 different neural networks that could successfully perform the task (Fig. S2A). The best neural network from this ensemble achieved a performance of 93.12%. Although this neural network correctly classified all probes, the performance score was not perfect due to slight deviations in the output (Fig. 3C).

In order to better understand how a neural network performed this task, we can characterize the flow of information across the agent-environment system. To this end, we decomposed the total information that the best neural network from the ensemble had about the cue into information uniquely transferred from the environment, uniquely transferred from the neural network, and redundantly from both, during the course of the task. During the cue stage, the environment was initially the unique source of information about the cue (Fig. 3D). As the neural network encoded the stimulus, the source became redundant. During the delay stage, the environment ceases to be a source of information. As the neural network had already encoded information about the cue, it becomes the unique source (Fig. 3E). Crucially, the neural network preserves this information throughout the delay stage. Finally, during the probe stage, the environment once again becomes a source, and therefore the source is redundant (Fig. 3F). Note that when the environment provides the probe stimulus it became the source of information about the cue. Since the neural network already contained information about the cue, the neural network and the environment both redundantly act as the source.

As explained previously, predictive information in this task arises from the relationship between cue and probe stimuli. Encoding information about the cue automatically results in encoding information about the probe (and vice versa). This is because knowing the cue significantly reduces uncertainty about the probe; the probe can only be one of two values given a cue. Predictive information that the neural network has about the probe and its sources is qualitatively similar to the information it has encoded about the cue (Fig. S3A). The neural network encodes information about the probe stimulus upon receiving the cue, and retains that predictive information during the delay stage. This is merely a consequence of encoding and retaining the cue. The entire ensemble of neural networks optimized to perform this task consistently exhibit this phenomenon of encoding information about the probe transferred uniquely from the cue stimulus (Fig. S3B) and is even robust to noise in the neural network (Fig. S5).

### Environmental regularities induces predictive information in any neural network

Since optimized neural networks encode information about the probe merely by encoding the cue, does any neural network that encodes the cue also encode information about the probe, and therefore have similar predictive information? In order to study this, we created 100 random neural networks and presented them with the same task. Although these neural networks were not able to perform the relational categorization task (Fig. S2B), they encoded similar amounts of total predictive information as the trained neural networks (Fig. 4A). Specifically, they encode the same amount of information about the probe during the cue stage (Fig. 4B). Furthermore, decomposing that information revealed that the information originated from the environmental stimulus and that the neural network dynamics had no role in its encoding of predictive information in both random and optimized neural networks (Fig. 4C). Thus, predictive information alone is not sufficient to distinguish neural networks optimized to perform specific tasks from random neural networks that are merely reflecting the information provided by the environment.

**Fig. 4.**
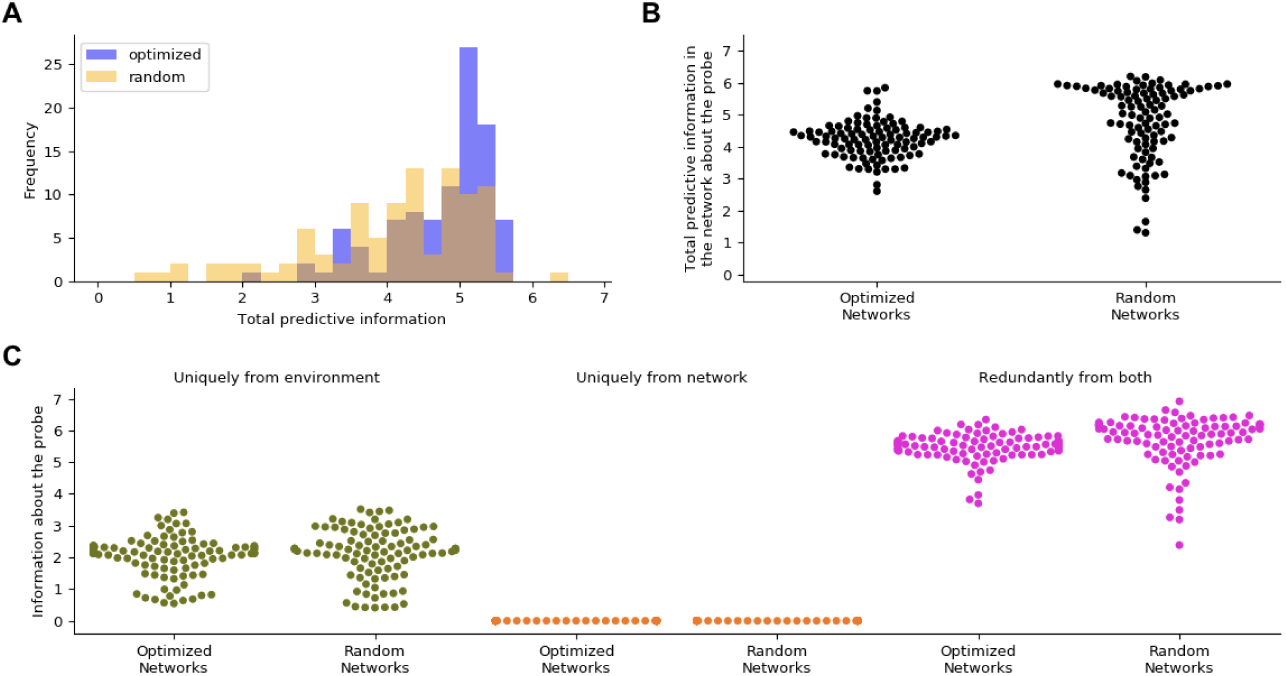
Comparison of predictive information sources in optimized and random neural networks. [A] Total predictive information estimated by averaging over the entire course of the task is similar in random and optimized neural networks. [B] Total predictive information about the probe averaged across the cue stage of the task, is the same in random and optimized neural networks. [C] Decomposition of that total predictive information showing that information about the probe in both random and optimized neural networks was from the environment (green), eventually becoming redundant as they both encoded the cue stimulus (pink). The neural network had no role to play in its encoding of predictive information about the probe during the cue stage (orange).

### Information decomposition distinguishes between random and optimized neural networks

Unlike CPG and PP that were distinguished based on having different information sources, random and optimized neural networks in the relational categorization task have the same information sources. Even under this condition, decomposing the total information across sources and unrolling over time helps distinguish them by revealing differences in the magnitude of information transferred from each source over time. Specifically, predictive information sourced by the neural network during the delay stage is markedly different between random and optimized neural networks. As discussed in the previous section, optimized neural networks preserve information about the cue (and hence predictive information about the probe) during the delay stage. In contrast, random neural networks tend to lose that information. As a consequence, the amount of unique information provided by the neural network at the end of the delay period is higher for the trained neural networks than for the random neural networks (Fig. 5A). This difference disappears when information is measured across time, and can only be observed by unrolling it over time.

**Fig. 5.**
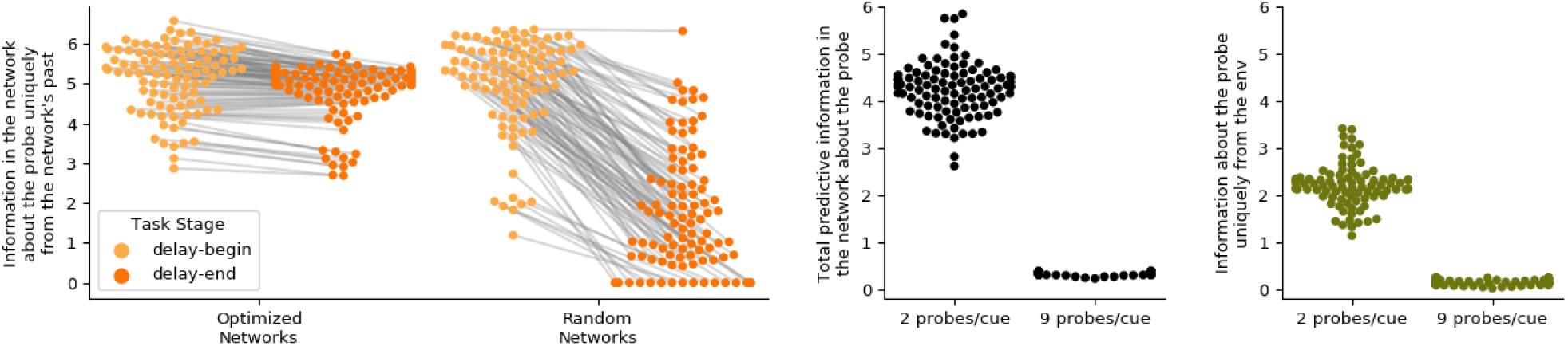
Influence of neural network and environmental properties on predictive Information [A] Both random and optimized neural networks have similar levels of information about the probe at the beginning of the delay stage (light orange), but unlike optimized neural networks, random neural networks lose that information by the end of the delay stage (dark orange). [B] Total predictive information in the optimized neural networks about the probe during the cue stage showed a significant drop upon changing environmental statistics from 2 probes/cue to 9 probes/cue. [C] Drop in total information show in B can be attributed to the drop in information uniquely from the environment about the probe in the 9 probes/cue setting.

### Statistics of the environment influences magnitude of predictive information

Encoding the cue results in encoding information about the probe in this task because of the relationship between them. How does changing this relationship impact predictive information in the neural networks? In order to study this, without changing the nature of the relational categorization task we merely changed the structure in the environment. This was achieved by modifying the task such that the probe could be one of 9 possible values for a given cue, rather than one of two possible values (Fig. S4B). Reduction in uncertainty about the probe’s value given the cue is now much less compared to the original environmental structure (Fig. S4D,E). This will be reflected in the information that the cue can provide about the probe. However, this came at no cost to performance because the neural networks were still encoding the cue just as well. The same ensemble of optimized neural networks were able to perform this task successfully without any more training (Fig. S4E). Information dynamics was then measured using data recorded under this 9-probe condition. Measuring the total information in the neural network during the cue stage about the probe revealed that there was significantly less information in the neural network in 9 probes per cue condition (Fig. 5B). The reduction in total predictive information can be wholly attributed to the reduction in information about the probe (Fig. 5C). Thus, differences in environmental structure can result in significantly different amounts of predictive information encoded in neural networks without any behavioral differences.

## Discussion

The study of predictive coding and its relevance to behavior has been studied from multiple perspectives in the literature with regards to the source of information: predictive information can be generated by the neural network (5, 6) and predictive information can be provided by the environment (7, 20). In this work, using computational models where the ground-truth about the source of information was known, we demonstrate that predictive information can originate from either the environment or the neural network or both, and that the source of information can dynamically change during the course of a behavior. In order to do this, we first presented a theoretical framework based on multivariate information theory that allows us to infer the source of predictive information and its dynamics. This involved decomposing the total information that neural networks encode about a future stimulus into information transferred uniquely from the neural network, uniquely from the environment and redundantly from both sources. We validated this framework using the CPG and PP models where information is known to originate from the neural network and the environment respectively. Second, using the more structured relational categorization task, we demonstrated that (a) amount of predictive information encoded in a neural network is not indicative of its performance; (b) the source of information about a future stimulus can change during the course of the task; and (c) the source of information about a future stimulus can change within the same task depending on the regularities of the environment. Thus, predictive information might be necessary but is not sufficient to explain the neural basis of a behavior. Decomposing information across sources and studying its dynamics over time takes us one step further in understanding the role of predictive information in a behavior.

The framework presented here for inferring the source of predictive information takes us beyond general correlations that information theoretic measured are known to capture by capturing the effects of perturbation on the neural system. Identifying the sources of predictive information requires that the system under study be perturbed. The presentation, removal or sudden change of a stimulus is a perturbation. This causes the system to break the redundant encoding observed in a steady-state. It is during such a perturbation that we can use partial information decomposition to determine the source of information in a coupled system. Once the neural network and the environment settle into the next steady-state after the transient due to the perturbation, information once again becomes redundant between them. Thus, through the combination of information decomposition, time-unrolling and perturbation we are able to infer the ground-truth causal influences in the models we have analyzed.

The framework presented here can be applied to experimental data across multiple scales. In fact, it can be applied to any time-series data spanning multiple trials corresponding to several perturbations from the steady state. However, in this work, we focus on open-loop systems. Specifically, we focus on agent-environment systems where the agent influences its environment or where the agent is influenced by the environment. Such an open-loop setup is typical in experiments in neuroscience, where the subject receives a stimulus, but does not have the ability to influence the future stimulus through their state or actions. In natural behavior, the agent and environment are in closed-loop interaction. The analysis of closed-loop systems introduces an added complexity. The regularities of the environment can be generated by the regularities of the neural network’s dynamics and vice-versa. As a result, the distribution of environmental stimuli and the distribution of the neural activity are dependent on each other, unlike the open-loop setup where one of them is independent of the other. As it is, the framework requires that one of the distributions be fixed across time in order to make fair comparisons of information at different time-points. Future work in this direction will involve extending the framework and designing the experimental setting that would allow us to infer the source of predictive information in a freely moving animal.

## Materials and Methods

In the agent-environment models used throughout this paper, the agents were modeled using dynamical recurrent neural networks. The parameters of the neural network were optimized using an evolutionary algorithm such that it was able perform the required task. In this section, we specify implementation details about the neural network model, the tasks, and the optimization algorithm.

### Neural network model

A Continuous-Time Recurrent Neural Network (CTRNN) was used as the model neural network (14, 15). The neural network consisted of three layers: the input layer which was connected by a set of feed-forward weights to the interneuron layer; the interneuron layer was a CTRNN which fed into the output layer; the output layer produced the output of the neural network which was given by a weighted combination of the interneurons’ output. The dynamics of each interneuron was governed based on state equations given by

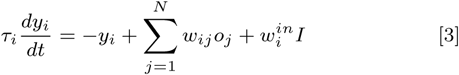

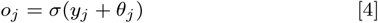

where *y*_*i*_ refers to the internal state of neuron *i*; *τ*_*i*_, the time-constant; *w*_*ij*_, the strength of connection from neuron *j* to neuron *i*; *o*_*j*_, the output of the neuron; *I*, the input and 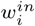, the weight from the input to the neuron. Based on the state of the neuron its output is given by equation 4, where *σ*() refers to the sigmoid activation function given by *σ*(*x*) = 1*/*(1 + *e*^−*x*^), and *θ*_*j*_ refers to the bias of neuron *j*. The output of the network at any time *t, O*(*t*), is estimated as a weighted sum of the outputs of each neuron (weights given by 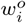), passed through a sigmoid function and scaled to be in the range [−1, 1].

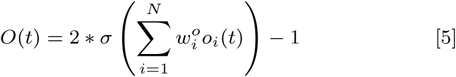

All neural networks described in this paper were made up of *N* = 3 neurons. The tunable parameters of such a model include the weights between the neurons (*w*_*ij*_), the input weights 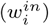, the output weights 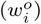, time-constants (*τ*_*i*_) and biases (*θ* _*ij*_) of each neuron. The model was simulated using Euler integration with a step-size of 0.02.

### CPG task

The neural network model described above is capable of intrinsically producing oscillations. To create Central Pattern Generators (CPGs), neural networks were optimized to produce oscillations from a range of initial conditions. The neural network was started at 100 different initial conditions by systematically setting the neuron outputs in the range [0, 1]. For each condition, the neural activity was recorded for 10 simulation seconds. The ability to generate oscillations was assessed by measuring the absolute difference in each neuron’s as well as the neural network’s output in consecutive time-steps across all time-points in a trial, and then across trials. The neural network’s output was fed to an environment governed by

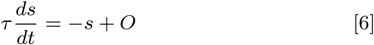

where *s* refers to the state of the environment, *τ* refers to its time-constant which was set to 0.5, and *O* refers to the output of the neural network given by equation 5.

### Relational categorization task

We adapted the relational categorization task to provide neural networks with structured stimuli (16, 17, 44). This task involves first providing the neural network with a cue stimulus in the range [3, 11] for 5 units of time. This is followed by a delay period when no stimulus is provided for 10 units of time. Finally, a probe stimulus that is of magnitude greater or less than the cue is provided for 5 units of time. The goal of the task is for the neural network to distinguish probes that were larger than the cue or smaller than the cue, by producing an output of +1 or − 1 respectively. In the first version of this task, the probe can take one of only two values, either *cue* + 1 or *cue* − 1. In the second version of the task, the probe can take any value in [3, 11]. While the goal of the task remains the same in both versions, the distribution of the probes given the cue, and therefore information that the cue gives about the probe is significantly different (Fig. S4). Performance of a neural network in this task was estimated by measuring absolute deviation of the network’s output from the desired output of +1 or − 1 during the probe stage. Time-averaged deviation was also averaged across all trials of cue-probe values, to obtain a score in the range [0, 1].

### Neural network optimization

Neural network models described previously were optimized to perform the relational categorization task using an evolutionary algorithm (46, 47). This optimization methodology involves instantiating a population of 100 random solutions that evolves over several generations to produce solutions capable of performing the task. A generation is defined as the process of creating a new population of solutions that has improved in “fitness” (task performance) from the last. Each solution, referred to as a genotype, is an *N* dimensional vector corresponding to the parameters to be optimized. The parameters were encoded to be in the range [0, 1] and scaled to produce the neural network that the genotype encoded. In each generation, the fitness of every genotype is evaluated and a new population is created using a fitness-based selection and mutation strategy as follows: The genotypes that perform in the top 1% were retained as is for the next generation. The rest of the individuals were created by selecting two genotypes preferentially in proportion to their fitness and combining them. To these offspring, Gaussian mutation noise with mean 0 and standard deviation 0.01 was added before being added to the population of genotypes for the next generation. After a fixed number of generations, the best individual in the population was selected as the representative solution from that optimization run. 100 such runs were conducted to obtain an ensemble of 100 neural network models that successfully performed each task. For the relational categorization task, optimization was carried out for 500 generations. In the case of the CPG task, at the end of 50 generations the optimization process was terminated and deemed successful if the best agent in the population reached a fitness of 30 or greater. This was repeated until 100 CPGs were produced. See supporting information (Figs. S1 and S2) for training curves, behavior of best optimized neural network, distribution of fitness of best models from 100 runs, and sample neural traces.

### Random neural networks

Matched random neural networks were created for the relational categorization task by shuffling the parameters of the optimized neural networks. All parameter groups, namely time-constants, input weights, recurrent weights, output weights, and biases were randomly shuffled within themselves rather than across groups. Thus, the ranges of parameters were preserved in each group but their associations with neurons were randomly shuffled.

### Measuring information transfer

To identify the source of information over time, information transfer measures were estimated independently at each time point. For any given time step, data for environmental stimulus at the previous time step, neural activity of previous time step, current neural activity, and stimulus at a future time step, was collected across multiple trials. Probability densities were estimated from this data using a kernel density estimation technique known as average shifted-histograms (48) with 7 shifted binnings of 100 bins along each dimension of the data space. These probability density estimates were then used to measure the redundant information terms in equation 2. Similar results were observed with 5 and 11 shifts and with 50 and 200 bins per dimension (Fig. S6). All information theoretic quantities were estimated from raw data using the *infotheory* package (49).

---

**Fig. S1.**
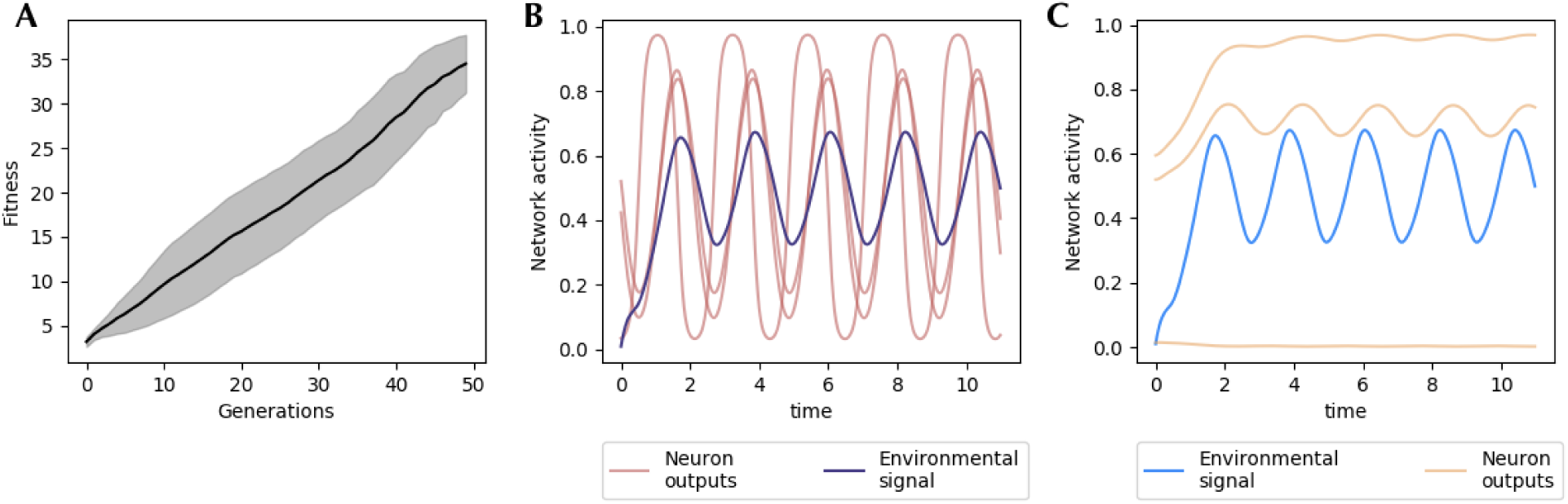
Optimization and neural traces of CPG and PP. [A] Fitness over time for 100 valid runs of optimizing a CPG model. Only runs that achieved a fitness greater than 30 were deemed valid. [B] Neural traces from one trial of the best CPG demonstrating that all neurons (red) as well as the neural network output (blue) oscillate. [C] Neural traces (orange) when the output from the CPG shown in panel B was fed to a random neural network in the PP condition demonstrating input driven oscillation in the random neural network.

**Fig. S2.**
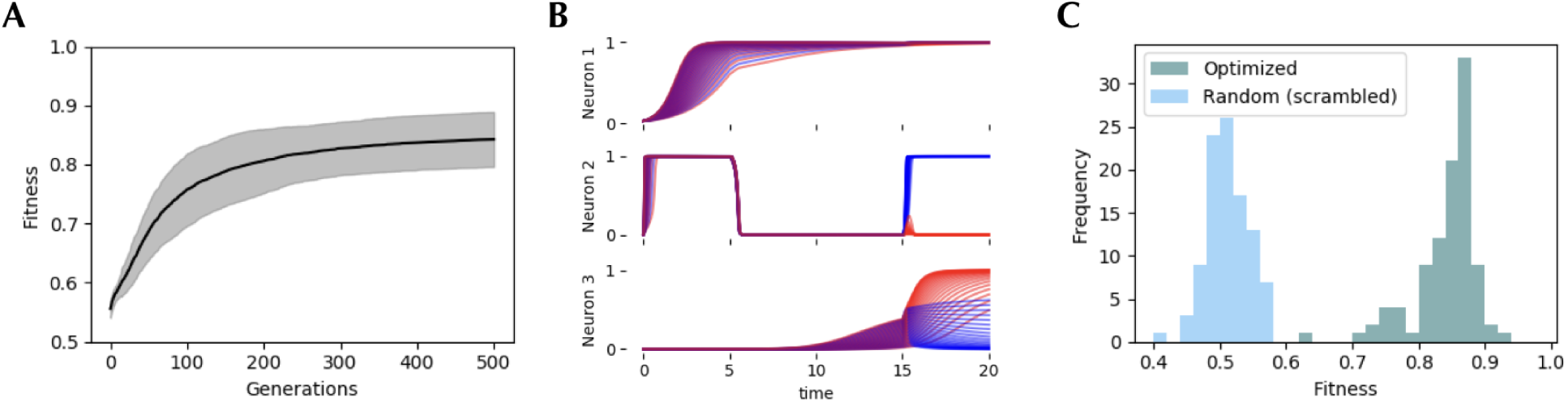
Optimizing neural networks to perform relational categorization. [A] 100 independent runs all converged to near-perfect performance with deviation from a perfect score only due to small deviations from expected output and not mis-categorization. [B] Neural activity in the CTRNN of the best optimized agent over 35 trials where probe was larger than the cue (red) and 35 trials where the probe was lesser than the cue (blue). [C] Neural networks whose weights and time-constants were scrambled lost their ability to perform the task.

**Fig. S3.**
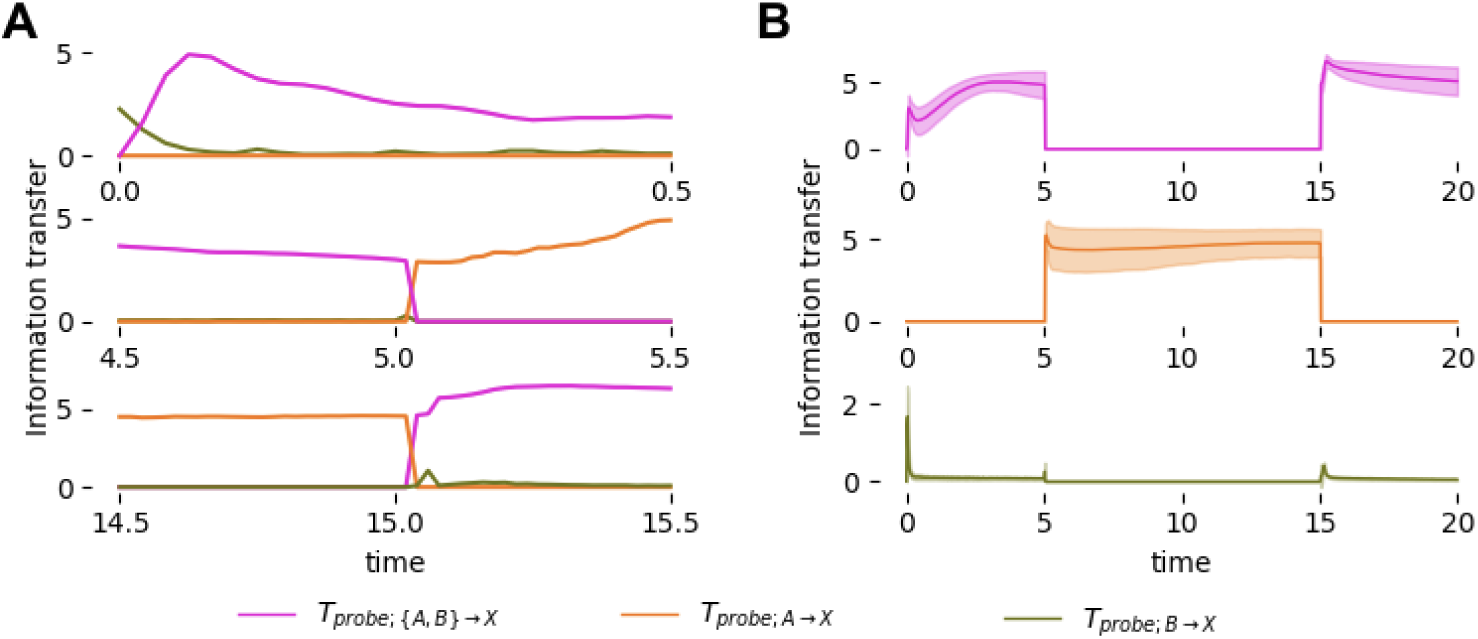
Predictive information source dynamics is consistent and similar with information about the probe. [A] At the start of the cue stage (top), information about the probe arrives from the environment (green) as the cue is provided, and becomes redundant as the cue is encoded (pink). Towards the start of the delay stage (middle), the neural network becomes the source of information about the probe (orange) as it retains information about the cue, and since the environment ceases to provide that information. As the probe is provided (bottom), the environment once again becomes a source of information in addition to the neural network and they are both redundantly sources of information (pink) [B] Predictive information source dynamics are consistent across all 100 optimized neural networks during all three stages of the task. Their mean value is shown in bold and the shaded region represents one standard deviation around it.

**Fig. S4.**
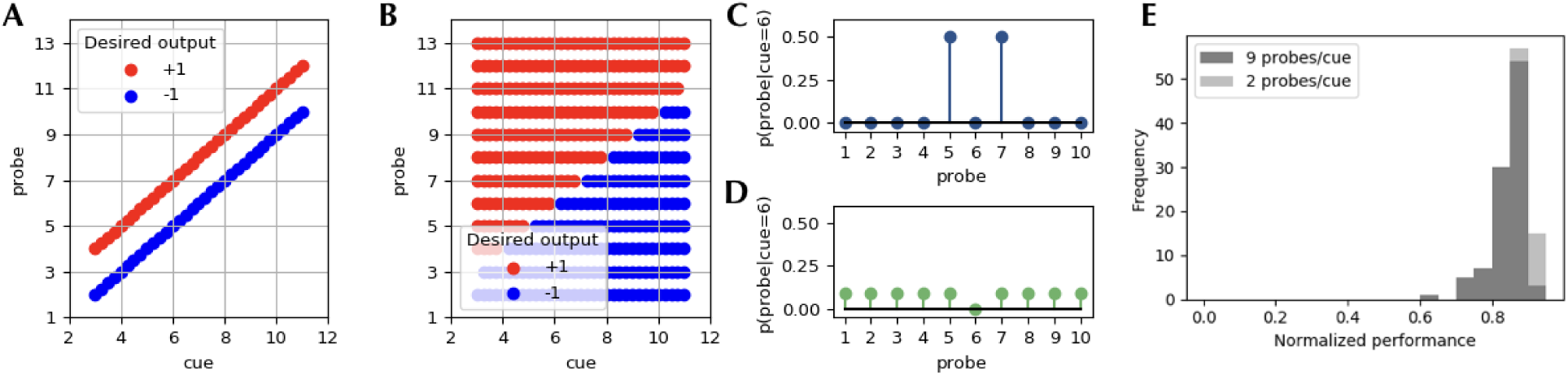
Different environmental structures within the relational categorization task [A] Relational categorization task with highly structured stimuli; for each cue probe is one of two possible values. [B] Relational categorization task with minimal structure in stimuli; probe can be one of 9 values for a given cue. [C] Conditional probability of probes given a cue for environmental structure shown in panel A, demonstrating the significant reduction in uncertainty of the probe given the cue. [D] Conditional probability of probe values given a cue under the environmental structure in panel B shows that probe values still have a nearly uniform distribution, and hence very less reduction in uncertainty. [E] Neural networks optimized to perform under the distribution shown in panel A perform just as well under the distribution shown in panel B.

**Fig. S5.**
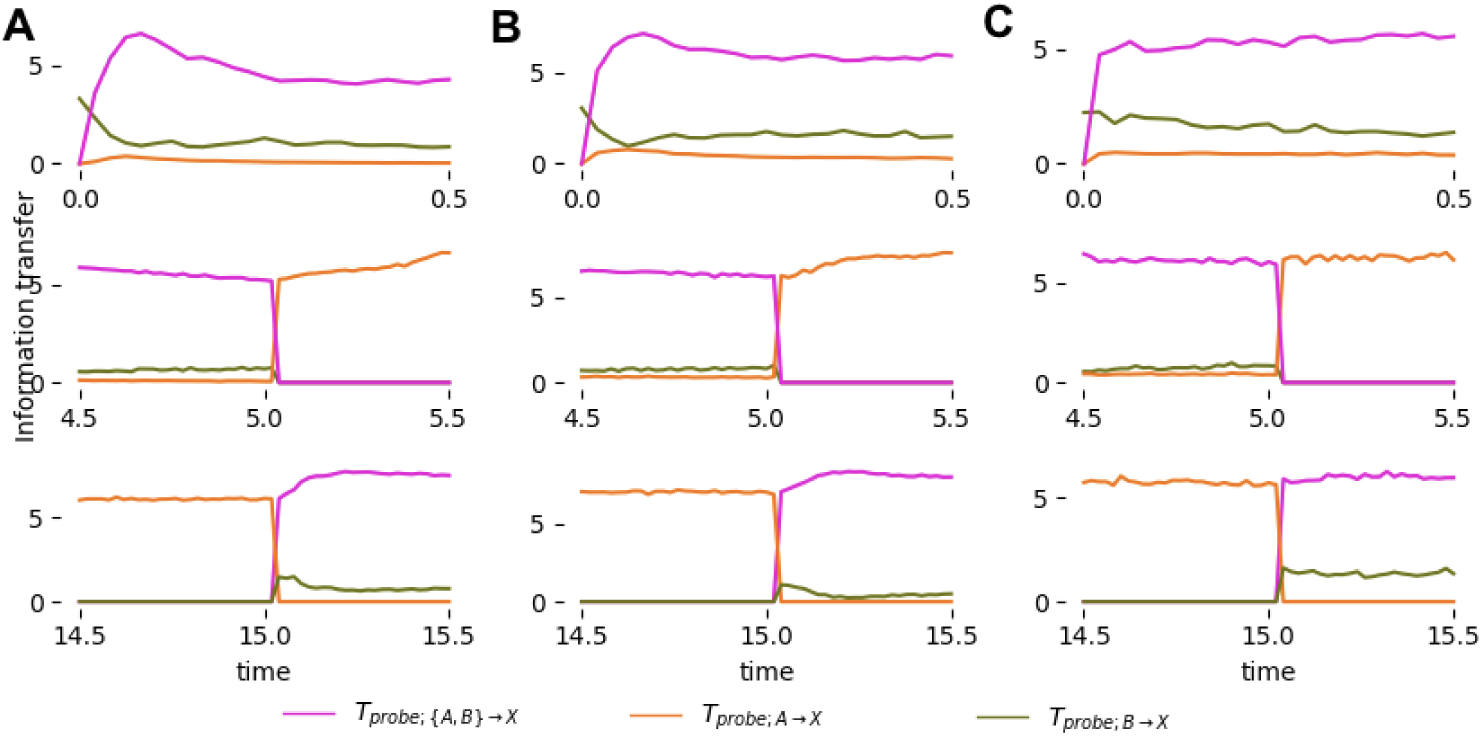
Inferring the source of predictive information is robust to zero-mean Gaussian noise with standard deviation [A] 0.01, [B] 0.05 and [C] 0.1. Results are qualitatively similar to results from fig. S3A for cue (top row), delay (middle row) and probe (bottom row) stages of the task.

**Fig. S6.**
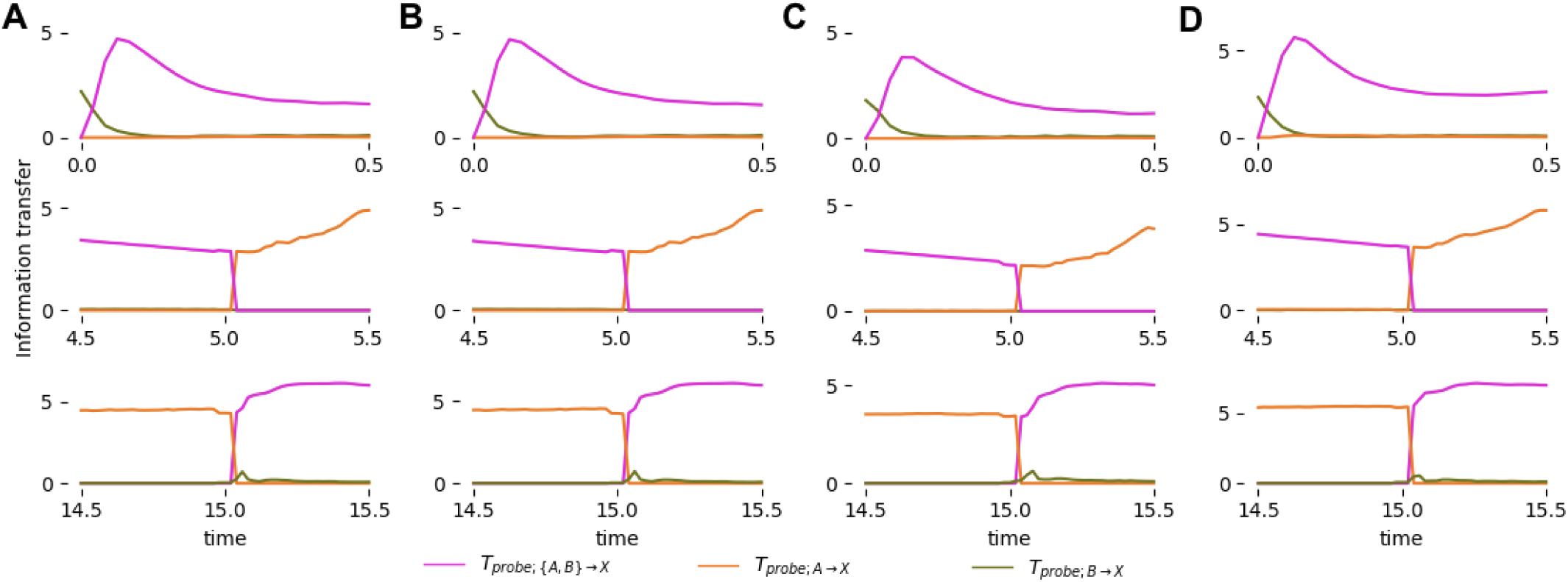
Inferring the source of predictive information with different binning and shifted-histograms. Results are qualitatively similar to results from fig. S3A after changing [A] number of shifted bins to 3 [b] number of shifted bins to 11 [C] number of bins per dimension to 50 and [D] number of bins per dimension to 200, for cue (top row), delay (middle row) and probe (bottom row) stages of the task.

